# Bumblebees retrieve only the ordinal ranking of foraging options when comparing memories obtained in distinct settings

**DOI:** 10.1101/2022.04.05.487177

**Authors:** Cwyn Solvi, Yonghe Zhou, Mark Roper, Yunxiao Feng, Li Sun, Rebecca Reid, Lars Chittka, Andrew Barron, Fei Peng

## Abstract

Are animals’ preferences determined by absolute memories for options (e.g., reward sizes) or by their remembered ranking (better/worse)? The only studies examining this question suggest humans and starlings utilize memories for both absolute and relative information. We show that bumblebees make decisions using only memories of ordinal comparisons. After learning to discriminate pairs of different flowers by sucrose concentration, bumblebees preferred flowers (in novel pairings) with 1) higher ranking over equal absolute reward, 2) higher ranking over higher absolute reward, and 3) identical qualitative ranking but different quantitative ranking equally. Despite these suboptimal choices, bumblebees found the optimal option in an ecologically relevant foraging task. Our results illuminate a divergent mechanism of learned preferences that may have arisen from adaptations to bumblebees’ natural environment.

## Introduction

What do animals remember about items out of context? For example, suppose we learn that different options (e.g., coffee shops) result in different reward outcomes (e.g., waiting time, quality) and later we’re presented with a choice between two previously encountered options which we’ve never experienced side-by-side. What types of values do we remember for those options now presented in a novel context? Do our memories of the subjective values for each option contain absolute information (e.g., delay to reward), remembered ranking (how they compared to previous alternatives), or a weighted combination of both?

In typical studies exploring the economic choices of animals including humans, subjects do not have to use distant memories of the options: they are presented with choices where the objective values (e.g. amount, cost, status) are concurrently visible and can be directly compared. Under such conditions, a wealth of research shows that animals’ choices can be influenced by the presence of additional options (Hunter and Daw, 2021; Spektor et al., 2021). An example of this phenomenon is frequently used in marketing: when given a choice between a $3 small popcorn and a $7 large popcorn most people choose the smaller cheaper option, but when a $6 medium popcorn is added, more people choose the large popcorn because it now seems like a good deal. Evidence of contextual effects like this on direct assessments has been found across the animal kingdom (e.g., humans and other primates (Berkowitsch et al., 2014; Parrish et al., 2015; Trueblood et al., 2013), bats (Hemingway et al., 2021), birds (Bateson, 2002; Morgan et al., 2012), frogs (Lea and Ryan, 2015), fish (Reding and Cummings, 2017), bees (Shafir et al., 2002), and worms (Iwanir et al., 2019)). However, little is known about the type and degree of information (absolute and/or relative) that is encoded in the *remembered* subjective values of options.

Only recently have investigations of absolute and relative information traversed into the realm of reinforcement learning, where value must be inferred from memories. An elegant study with starlings (Pompilio and Kacelnik, 2010) and a series of impressive studies with humans (Bavard et al., 2021, 2018; Klein et al., 2017) demonstrated that both absolute memories and remembered ranking are combined in particular ways to give rise to these animals’ preferences, i.e. use of both absolute and ranking information is required to explain both species’ behaviors. So far, however, no other species has been investigated for the roles of absolute and relative information in learned preferences. Here, we examine this in bumblebees, an invertebrate and a key model for examining the economy of decision-making outside of humans (REAL, 1996).

## Results

If bumblebees encode and retrieve memories for absolute values, their preference for a particular option should not depend on the context in which that option was learned (Padoa-Schioppa and Assad, 2008). To test this idea, we conducted experiment 1 with a multi-contextual design where certain flowers had the same quality of reward but different ranking. Bees were first trained (individually in all experiments) on two different pairs of colored flowers (A & B; C & D; Figure 1A, Figure 1—figure supplement 1, and Figure 1—figure supplement 2). The sucrose concentrations of the different pairs of flowers were chosen according to Weber’s Law (Akre and Johnsen, 2014) to represent the same perceived difference, i.e. A:B = 45%:30% had the same contrast of incentives as C:D = 30%:20% (Figure 1A). Following sequential training in both contexts (training sequence and color-reward contingency was counter-balanced across bees; Figure 1—figure supplement 3 and Figure 1—figure supplement 4), we tested each bee’s preference between flower types B and C, which had the same reward quality (30%) during training. Therefore, if bumblebees remembered absolute values for options, their preference between B and C in the test should be identical. Any significant deviation would suggest a contextual effect due to B and C having different rankings during training (B < A while C > D). In an unrewarding test with B and C flowers, bees significantly preferred C over B (GLM: 95% CI = [0.08 0.38], N = 40, P = 4.00e-3; Figure 1B and Figure 1—figure supplement 3). Although these results suggest that bees store memories for the relative ranking of flowers, they still may encode absolute information (Table 1) and therefore we carried out additional experiments.

**Figure 1.**
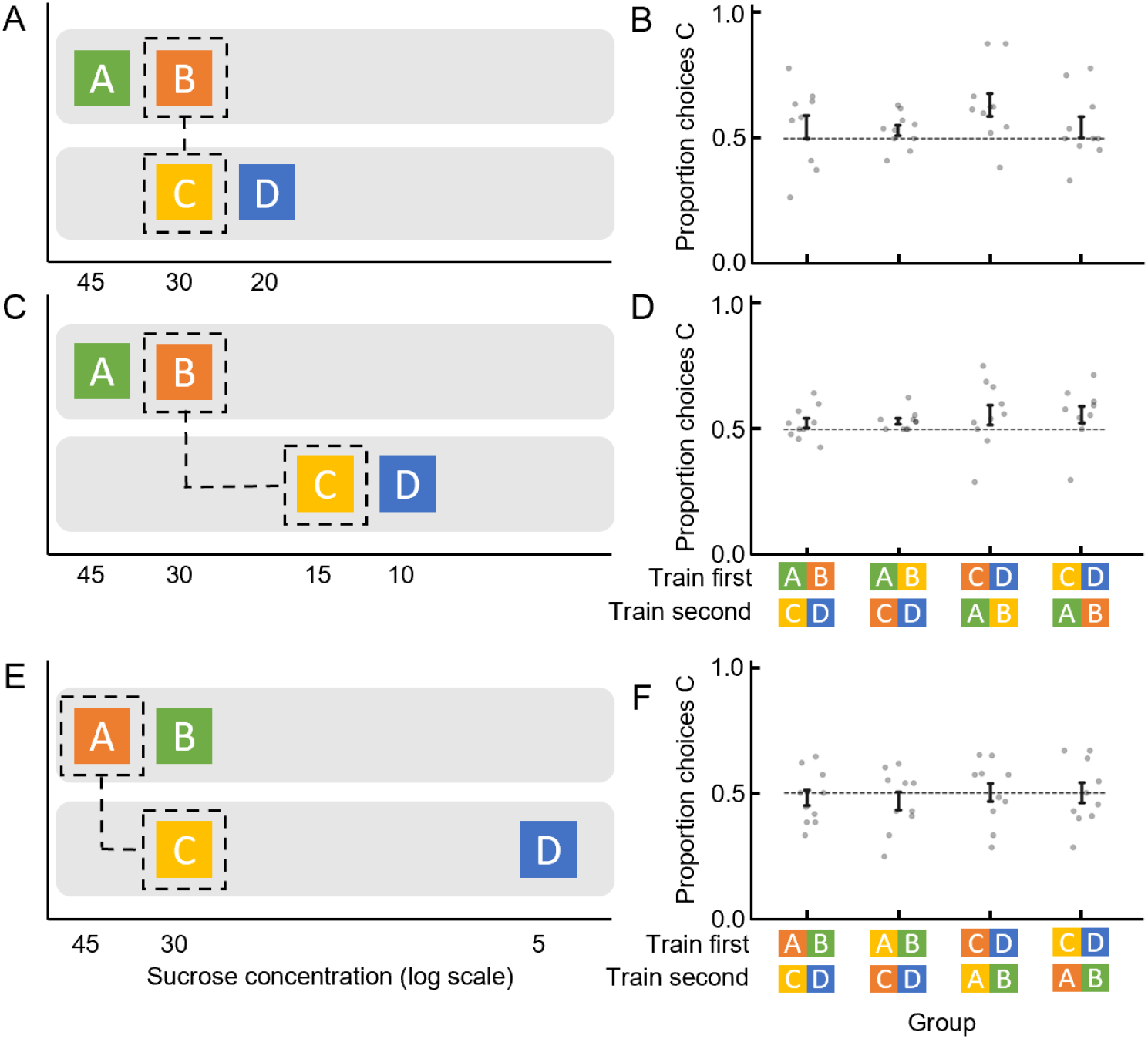
Bumblebees make decisions based on ordinal comparisons. **(A), (C), (E).** The corresponding sucrose concentration of each stimulus is displayed on a log scale to visually represent their relative differences according to Weber’s Law (Akre and Johnsen, 2014). Training sessions are indicated by separate grey backgrounds and the test options in each experiment are indicated with dashed lines. **(B), (D), (F).** Test results for each experiment. Groups indicate different counter-balanced training sequence and color-reward contingency (see Figure 1—figure supplement 3 for more details). Dotted horizontal lines indicate chance performance. Vertical lines indicate mean ± s.e.m.

**Table 1.** Predictions for various decision strategies for bumblebees’ flower preferences in experiments 1, 2, and 3. The left column lists the different categories of decision strategies. The middle three columns show the predicted results of each strategy for each experiment. The right column shows whether the predictions of each strategy match the behavior of bumblebees in all experiments. A lexicographic combination strategy is where choices are determined by a difference in a priority dimension (e.g. ranking), but if options are equal in that dimension then choices rely on a secondary dimension (e.g. absolute memory). A non-lexicographic combination strategy is where differences in either dimension can be used to make a choice. Value by association entails one option acquiring a higher value because it was experienced in a richer environment (Pompilio and Kacelnik, 2010).

| Strategy | Expected preference |  |  | Matches bees' behaviour |
| --- | --- | --- | --- | --- |
|  | Experiment 1<br>B(30%) vs C(30%) | Experiment 2<br>B(30%) vs C(15%) | Experiment 3<br>A(45%) vs C(30%) |  |
| Absolute memory | Indifferent | B | A | × |
| Remembered ranking | C | C | Indifferent | ✓ |
| Lexicographic combination—Absolute memory priority | C | B | A | × |
| Lexicographic combination—Ranking priority | C | C | A | × |
| Non-lexicographic combination | C | B/C/Indifferent | A | × |
| Value by association | B | B | A | × |

The following figure supplements are available for figure 1.

**Figure supplement 1**. General setup for experiments 1, 2, and 3.

**Figure supplement 2**. Specifications of the colours used for all the experiments.

**Figure supplement 3.** Counterbalanced colour sets used in experiments 1-4.

**Figure supplement 4.** Bees can discriminate flowers of different colours and sucrose concentration.

Bees may still store and use memories for both absolute and relative information in some combined fashion. To test this, in experiment 2 we set the reward quality of flowers B and C to be different (Figure 1C and Figure 1—figure supplement 3). The sucrose concentration of flowers was A > B >> C > D (45%, 30%, 15%, 10%). After training in both contexts, bees’ preference between B and C flowers was tested. If bumblebees have memories for both ranking and absolute information, they should either prefer B over C because it is much higher in quality, or this value difference would cancel out the ranking difference (B < A while C > D) and their preference for B and C should be equal. On the other hand, a preference for the much lower reward quality flower C would indicate they have access to only ranking information. Similar to experiment 2, the result of the unrewarding test showed that bees preferred C (GLM: 95% CI = [0.07 0.28], N = 40, P = 3.30e-03; Figure 1D).

Note that a preference for C over B in experiments 1 and 2 may have resulted from remembered ranking or a distorted absolute memory for flower type B due to the presence of and comparison with A during training. Evidence for the latter would manifest as a difference in the preference for C over B in experiment 2 compared to in experiment 1, whereas remembered ranking would predict a similar preference. There was no difference in preference for C across experiments 1 and 2 (GLM: 95% CI = [−0.24 0.13], N = 80, P = 0.55; Figure 1B and D). These results further support the notion that bumblebees’ choices in novel contexts are based on remembered ranking. However, one more experiment was necessary to determine if absolute memories may still be used by bumblebees (Table 1).

If bumblebees are relying purely on remembered ranking, then their memories for options should only be ordinal (Vlaev et al., 2011). That is, they can tell one option is better than another but cannot tell how much better. If so, there should be no indication that bumblebees can compare options in quantitative terms. To test this, in experiment 3 we made the reward contrast ratios within the two contexts very different ([A:B] = [1.5:1] = 45%:30% and [C:D] = [6:1] = 30%:5%; A > B = C >> D; Figure 1E). Here, testing bees’ preference between A and C flowers can answer our question. If bees encode memories for relative differences in quantitative terms, C should be preferred over A, as C had a higher relative value than A. Alternatively, if bees only remember the ordinal rank of options, preference for A and C should be the same because they have the same rank within their training contexts. In the unrewarding test, bees chose A and C equally (GLM: N = 40, 95% CI = [−0.16, 0.12], N = 40, P = 0.79; Figure 1F), suggesting bumblebees store memories for only the qualitative difference between options. Further, bumblebees’ equal preference for the equally ranked A and C flowers, despite their difference in sugar concentration, demonstrates absolute memories were not used (Table 1) and, in combination with experiments 1 and 2, confirms that bumblebees only use remembered ranking.

The results of our experiments highlight the potential suboptimality of using only remembered ranking. In experiment 4, we set out to test how this strategy performs in a semi-realistic foraging situation. In a multi-contextual setting with a divided arena, bees learned to forage from flowers of four different colors (Figure 2A and Figure 1— figure supplement 3). Importantly, bees could only see flowers of two colors at any one time, i.e. could only see A and B (45% and 30%) flowers from one side of the arena and could only see C and D (30% and 20%) flowers from the other side (Figure 2A). This setup mimicked a natural foraging situation with two distally separated patches of different flowers with different nectar concentrations. In the unrewarding test with all four options available, bees preferred option A (GLM: 95% CI = [0.94, 1.68], N = 20, P = 1.21e-06; Figure 2B), indicating that despite being susceptible to suboptimal outcomes in lab-created situations, using only memories of options’ ordinal ranking may be evolutionarily rational, i.e. can still lead to optimal choices within an ecologically relevant scenario.

**Figure 2.**
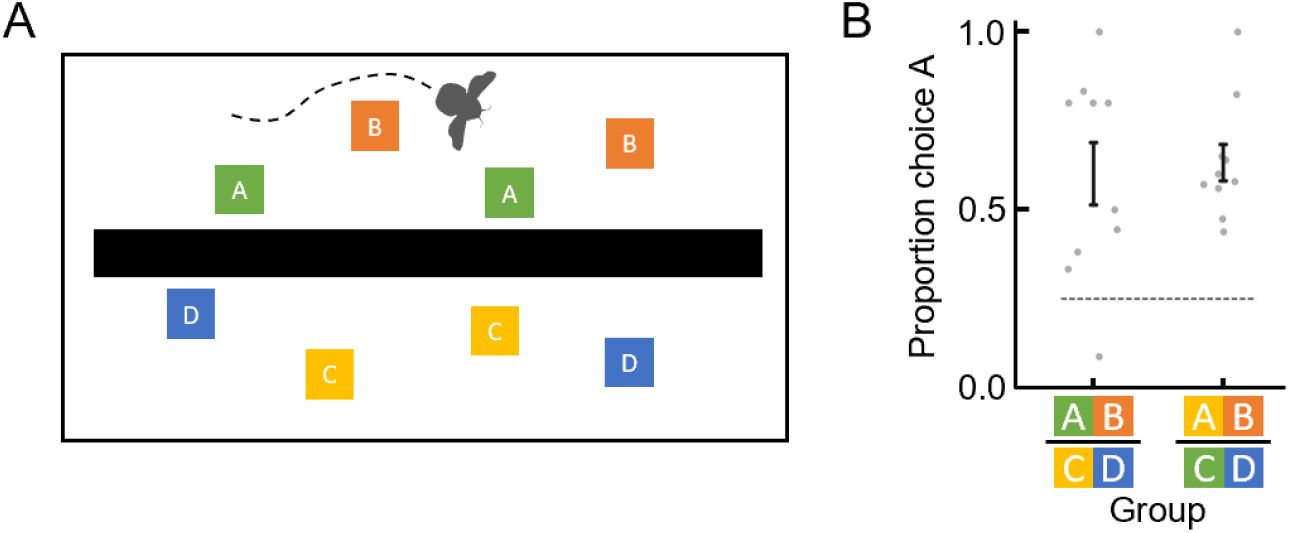
A suboptimal strategy can result in optimal foraging outcomes. **(A).** Top view of setup for experiment 4 where a wall separated flowers such that the bee could not see both groups of flowers at the same time. **(B).** Results of the unrewarding test in experiment 4. Groups indicate different color-reward contingency for bees (Figure 1—figure supplement 3). Dotted horizontal lines indicate chance performance. Vertical lines indicate mean ± s.e.m.

## Discussion

Why then would bumblebees have evolved to use only memories for ordinal comparisons while humans and starlings evolved to use both absolute and ranking information? Breadth of diet has been suggested to play a role in the evolution of cognition (Hemingway et al., 2017; MacLean et al., 2014; Simons and Tibbetts, 2019). Humans and starlings forage on a range of different foods, whereas adult bumblebees feed almost exclusively on nectar from flowers. Perhaps a varied diet, whereby one might need a common currency across vastly different food types (Chib et al., 2009; Levy and Glimcher, 2012) may have forced some animals to retain and use absolute memories, whereas a limited diet and limited dimensionality of reward might have favored ordinal relationships alone.

There is no reason yet to suggest a bee’s brain lacks the neural substrates to compute and store absolute memories for options. The basic information signal transferred between neurons is quantitatively expressed as frequency and pattern of action potentials. These signals could theoretically be used by higher centers of the bumblebee’s brain to encode an option’s utility as a cardinal (absolute) value (Miriyala et al., 2018; Schultz, 2015). However, our results suggest that bumblebees do not access or utilize this information when it would be of benefit in novel contexts.

Whatever the ultimate and proximate causes of bumblebees’ ordinal-only memories for options’ values, our findings demonstrate a fundamental difference in the mechanisms underlying learned preferences between those of bumblebees and those of starlings and humans.

## Materials and Methods

### Animals and setup

Bumblebee colonies were obtained from the Chinese branch of the Biobest Group (Biobest Belgium N.V., Westerlo, Belgium) and housed in wooden nest boxes (28 cm x 16 cm x 11 cm). Although there are no current requirements regarding insect care and use in research, experimental design and procedures were guided by the 3Rs principles (Russell and Burch, 1959). The behavioral tests were non-invasive and the types of manipulations used (sucrose, water) are all experienced by bumblebees during their natural foraging life in the wild. The bumblebees were cared for on a daily basis by trained and competent staff, which included routine monitoring of welfare and provision of correct and adequate food during the experimental period. A foraging arena (40 cm x 59 cm x 41 cm; Figure 1—figure supplement 1) was connected to the nest boxes via an acrylic tunnel with sliding doors, allowing experimenters to control bees’ access to the arena. Individual bees were marked with number tags (Opalithplättchen, Warnholz & Bienenvoigt, Ellerau, Germany), which were super-glued to the bees’ thorax. To ensure the sucrose concentration used in experiments was motivating, colonies were fed outside of experiments with 5-15% sucrose solution (w/w). They were also provided with ~3 g pollen every day. Illumination was provided by daylight fluorescent tubes (MASTER TL-D 90 DeLuxe 36 W/965, Philips, Eindhoven, the Netherlands) and near-ultraviolet fluorescent tubes (TL-D 36 W BLB, Philips) with high-frequency electronic ballasts (EB-Ci 1-2 36W/1-4 18W, 42-60 kHz, Philips), to generate a flicker frequency beyond the bumblebee’s flicker-fusion frequency. Coloured acrylic squares (25 mm x 25 mm x 5 mm) set on top of opaque glass cylinders as (artificial) flowers were placed in the arena with a different random spatial arrangement each trial. The spectral reflectance (Figure 1—figure supplement 2A) of all flower colours used in experiments was measured with a wavelength range of 300 – 700 nm and with 1 nm increments, using a spectrophotometer (Ocean Optics USB 2000+; Shanghai, China) and a deuterium/halogen light source. The perceptual positions of the colours in the bee colour hexagon space (Figure 1—figure supplement 2B) were calculated using the spectral reflectance measurements and the published *Bombus terrestris* spectral sensitivity functions of their three photoreceptors (Chittka, 1992).

### Experimental protocol

All bees involved were individually pre-trained on eight transparent artificial flowers, and were allowed to collect a full crop of sucrose solution from those flowers. Once a bee had successfully foraged for at least three consecutive bouts, she was moved to the training phase of an experiment.

#### Experiments 1, 2 and 3

Bees (N = 40 for each experiment) were trained to forage from eight flowers, four of each of two colours (Figure 1—figure supplement 1 and Figure 1—figure supplement 2). In experiment 1, one group of bees (n = 10) learned (individually) that blue flowers contained 45% sucrose solution and that yellow flowers contained 30% sucrose solution. Once training on these two flower types was complete (more detail below) this group of bees learned (individually) that orange flowers contained 30% sucrose solution and that green flowers contained 20% sucrose solution. Training for bees in experiments 2 and 3 was the same except the sucrose concentrations were different. Training sequences and colour combinations were equally counterbalanced across bees in each experiment (Figure 1—figure supplement 3). Flower colours and sequences for all experiments are listed in Figure 1—figure supplement 3. Initially during training all bees would land and drink from both flowers. As bees learned the colour-reward contingency, they began to land on the lower rewarded flowers but did not drink. Once this behaviour was observed, during subsequent bouts we removed and replaced one higher-ranking flower with one lower-ranking flower. By doing so, and because none of the flowers were refilled during training bouts, bees would again visit the lower-ranking flowers and drink the sugar water droplet. This method helped us to ensure that bees experienced drinking sucrose solution from both options an equal number of times. Training was completed when a bee collected approximately 50 of the 20μl aliquots of sugar reward from flowers of both colours. Note that we verified, in our setup, that bees could learn to discriminate between flowers of different colours and sucrose concentrations (Figure 1—figure supplement 4).

#### Experiment 4

Bees (N= 20) were trained and tested similarly to the other experiments except that the two sets of flowers were in the arena at the same time and accessible to the bee during each bout, but they were separated by an opaque wall to prevent the bees from seeing both sets simultaneously. The wall extended to cover most but not the full length of the arena, and there was space at the entrance and the far-end of the arena so that bees could freely choose to visit flowers on either side of the wall. No matter where the bee was in the arena, she could not see both pairs of flowers at the same time. In addition, flowers were refilled after the bee emptied the flower and began drinking on another. Each group (n = 10) was trained with either the yellow or green flower as the high concentration flower. The training was deemed complete when either a bee performed 200 landings or when a bee only landed on the mostly rewarded option for three consecutive bouts. After training, the bees were individually tested with all four flowers (two of each type) presented in the same arena without the opaque barrier.

### Statistical analyses

R v.3.6.1 was used to perform all generalised linear models (GLM) with quasibinomial distribution and logit link function. For all models, the response variable was the proportion of choices for the flower that had been associated with high concentration sugar. Fixed factors for models for results of experiments 1-4 were i) colony of each bee, ii) flower colour, and iii) training sequence (whether the higher concentration sugar reward was used during the first or second training session) and none were found to have a significant effect on colour preference during the tests in any experiment. When comparing preference for flower C across experiments 1 and 2, experiment was set as a fixed factor. Significance of fixed effects was tested by comparing full and null models using likelihood ratio tests.

## Supplementary Figures

**Figure 1 – figure supplement 1.**
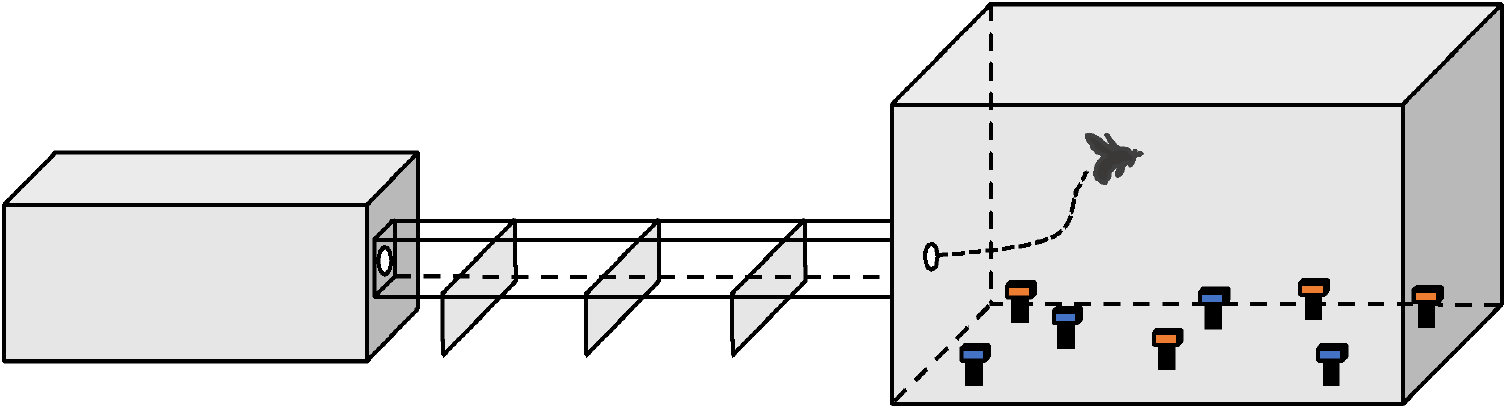
General setup for experiments 1, 2, and 3. In each experiment, artificial flowers of paired colours were horizontally presented in the training and testing phase.

**Figure 1 – figure supplement 2.**
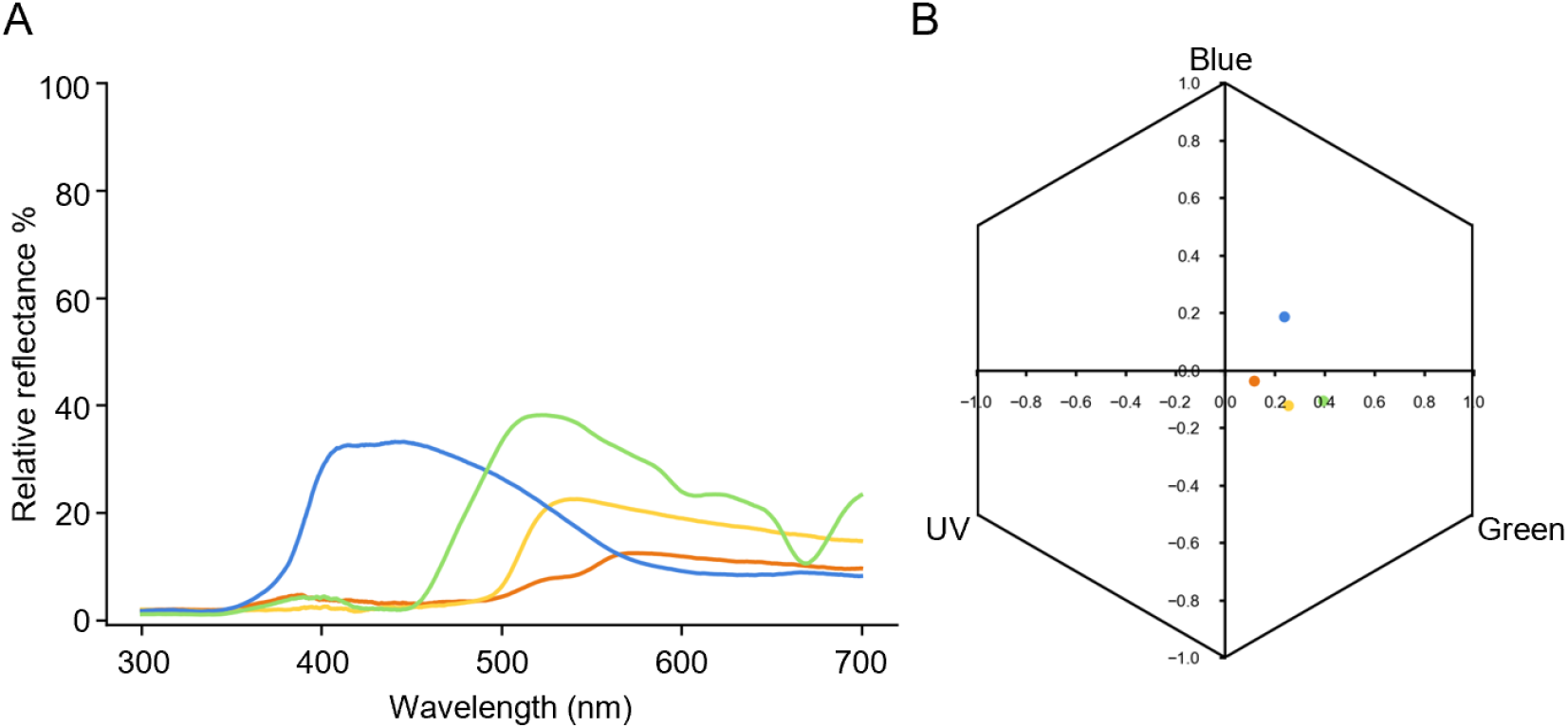
Specifications of the colours used for all the experiments. A. Spectral reflectance plot of dark blue, light blue, orange, yellow and green colours used. B. Loci of colours in the hexagonal bee colour space, determined by the responses each colour elicits on the bee’s ultraviolet, blue, and green photoreceptors (Chittka, 1992).

**Figure 1 – figure supplement 3.**
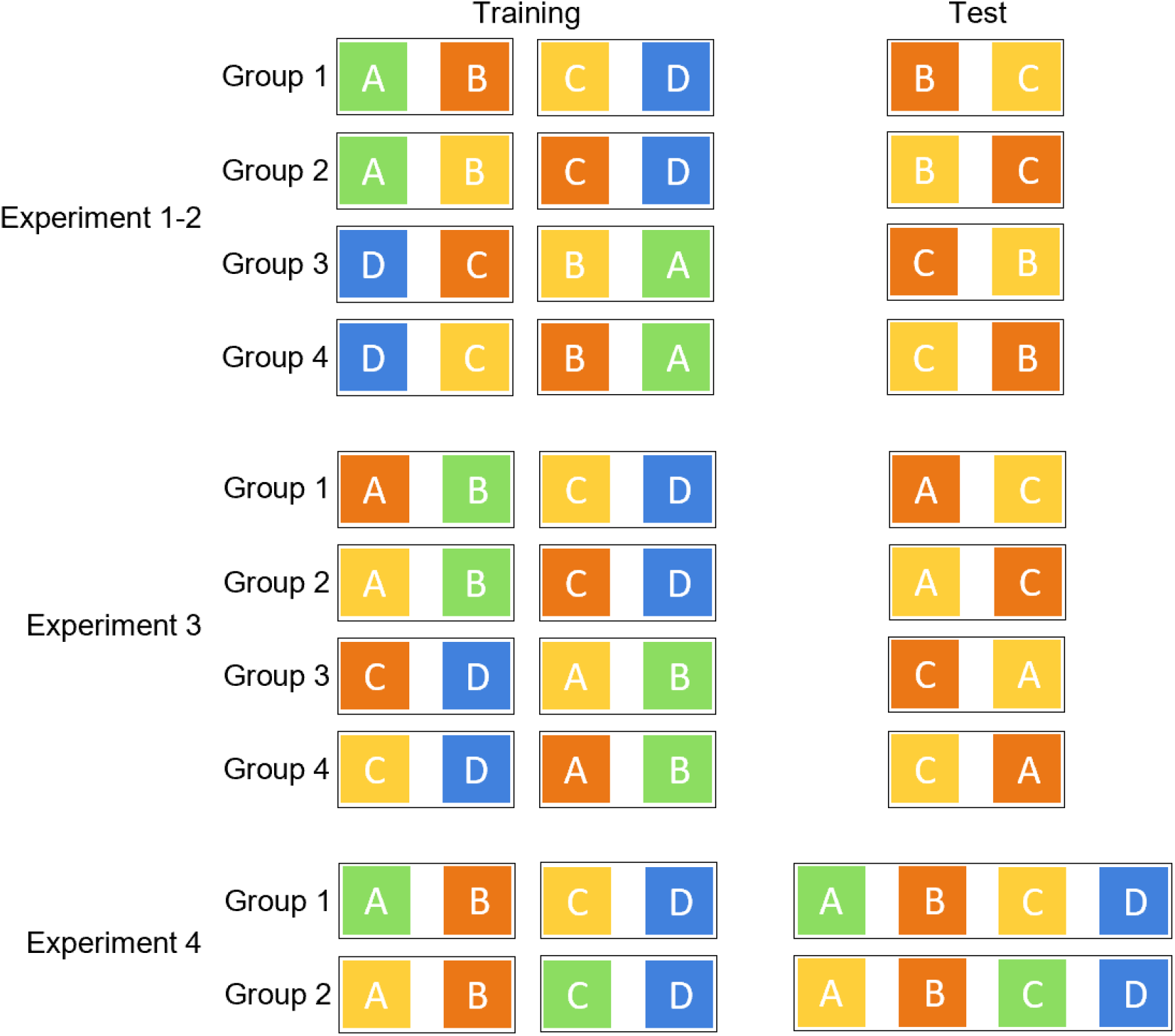
Counterbalanced colour sets used in experiments 1-4. Groups of bees were trained and tested with counterbalanced colour sets and training sequences in each of the different experiments.

**Figure 1 – figure supplement 4.**
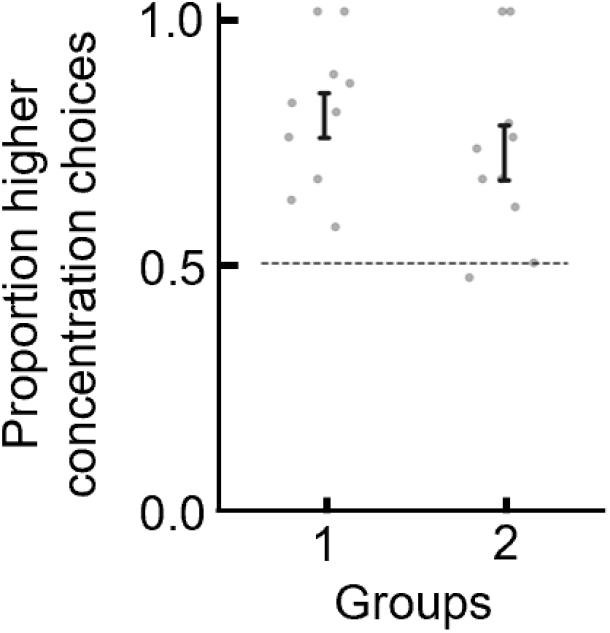
Bees can discriminate flowers of different colours and sucrose concentration. To ensure that bees were able to learn to discriminate between two differently coloured flowers in our setup, we examined bees’ preference between two flower types, after having been trained individually on these flowers. One group of bees (n = 10) learned that blue flowers contained 45% sucrose solution and that yellow flowers contained 30% sucrose solution. Another group of bees (n = 10) learned the counter-balanced colour-reward contingency. During a subsequent unrewarded test (all flowers with unrewarding water), bees showed a clear preference for (landed more often on) the flowers that had been associated with the higher reward during training (GLM: N = 20, 95% CI = [0.64 1.37], P = 3.39e-05). These results show that in our setup bumblebees were able to easily learn to discriminate the different flower colours used in our experiments, and do so via the different sucrose concentrations associated with each flower type. Groups indicate different counterbalanced colour-reward contingency for bees. Dotted horizontal lines indicate chance performance. Vertical lines indicate mean ± s.e.m.

